# Stimulation of saliva affects the release of aroma in wine; a study of microbiota, biochemistry and participant origin

**DOI:** 10.1101/2024.02.25.581677

**Authors:** Xinwei Ruan, Yipeng Chen, Aafreen Chauhan, Kate S. Howell

## Abstract

Saliva influences the release of aroma in the oral cavity. The composition of human saliva varies depending on stimulation and host’s origin; however, the compositional differences of saliva and their influences on aroma release have not been fully evaluated. In this study, we recruited 30 healthy adults (15 Australians and 15 Chinese) and collected saliva samples at three stages: before, during, and after stimulation. Salivary samples were characterised by flow rate, total protein concentration, esterase activity, microbiome composition by full-length 16S rRNA gene sequencing, and the ability to release aroma from wine by headspace solid-phase microextraction gas chromatography–mass spectrometry (HS-SPME-GC-MS). Differences in salivary composition and specific wine volatiles were found between Australian and Chinese participants, and amongst the three stimulation stages. Significant correlations between the relative abundance of 3 bacterial species and 10 wine volatiles were observed. Our results confirm the influence of participant’s geographic origin and stimulation on salivary composition, highlighting the role of salivary components, especially salivary bacteria, on the release of aroma from wine.

## 1 Introduction

Saliva plays an important role in aroma release from foods and beverages ^1–3^. When mixing with saliva in oral cavity, volatile compounds are released from food matrix and then travel to olfactory receptors via a retronasal route. Physiochemically, saliva has a dilution effect on aroma due to continuous salivation ^4^. Saliva contains salt, sugar and the proteinaceous fraction of mucin, which may influence how volatiles partition in a solution ^5^. Further biochemical effects by degrading odorants or releasing aroma from aroma precursors have been reported ^6–8^. Two critical components of saliva, proteins and microbiota, are implicated with release or formation of aroma compounds and thus affect aroma release from food

Proteins are able to bind, trap and interact with volatile compounds, thereby influencing the aroma release ^9^. Two abundant proteins in whole human saliva, mucin and α-amylase, modify release of aroma compounds through hydrophobic interactions ^10^. Pérez-Jiménez, Rocha-Alcubilla and Pozo-Bayón ^11^ demonstrated the efficacy of esterase in human saliva to degrade esters by comparing whole saliva with enzymatically inhibited saliva. Large inter-individual variations have been observed in the total salivary esterase activity ^12^. The chemical structure of esters also influences the extent of enzymatic degradation^6^. In addition, the effect of salivary enzymatic reactions on aldehydes ^7^ and thiols ^6^ has also been demonstrated by comparing the aroma release ability of thermal-treated saliva with untreated saliva.

In contrast to salivary proteins, the role of oral bacteria in aroma release is not well characterised. The oral microbiota contribute to the ability of human saliva to hydrolyse odourless grape glycosides ^8, 13^. A previous study ^14^ identified the differences in salivary microbial profile between obese and normal-weight individuals. However, no association was established between the differences of these two groups in aroma release of wine and the microbial profiles. In recent years, the advances in culture-independent methods for oral microbiome identification have deepened our understanding of microorganisms in the oral cavity ^15^ and reveal the salivary microbial composition with a stable core bacterial community shared by geographically diverse populations ^16^. Increased accuracy of taxonomic identification would facilitate to study the role of salivary bacteria in aroma release.

Salivary protein and microbiota are likely to be dynamic during oral processing. Saliva production switches between stimulated and unstimulated depending on the presence of stimuli. During stimulation evoked by taste or mastication, the saliva secretion volume increases ^17^. More than 90% of saliva comes from three pairs of major salivary glands, including the submandibular, sublingual, and parotid glands. The saliva produced by these three glands differs in composition and function because of the cells that constitute the glands ^18^. For example, the parotid glands secrete watery fluids since they exclusively consist of serous acinar cells, while the sublingual glands produce viscous saliva rich in mucins and other glycoproteins due to the high proportion of mucous acinar cells ^19^. Therefore, the salivary composition varies depending on the contribution of different glands with or without stimulation ^20^. It has been documented that parotid glands have a higher secretory rate under stimulation than in unstimulated conditions. Meanwhile, the submandibular and sublingual glands produce more saliva under unstimulated conditions ^21^.

Recently, the published literature has offered contradictory findings on whether the microbial diversity in saliva is different between unstimulated and stimulated status ^22–24^. Specifically, the alpha-and beta-diversity of salivary microbiome were reported as significantly different by Gomar-Vercher, Simón-Soro, Montiel-Company, Almerich-Silla and Mira ^22^. In contrast, the other two studies observed no significance ^23, 24^. Significant differences have also been observed by previous studies between saliva collected before and during stimulation in various salivary biochemical parameters, including protein concentration ^25^, antioxidant activity ^25^, esterase activity ^26^. Meanwhile, the composition of saliva collected after stimulation by food or other stimuli have not been characterised. Since the after-odour and after-taste generated after the swallowing or expectoration of food is also considered an essential sensory parameter ^27, 28^, studying the salivary composition after stimulation may give insight into understanding this process.

Meanwhile, the microbiota and type of protein in saliva varies among populations and is also shaped by the host characteristics, such as age ^29^, gender ^30^, body mass index (BMI) ^31^ and other lifestyle factors^32^. The correlation of a host’s ethnicity on the bacterial salivary composition has received considerable critical attention. Mason, Nagaraja, Camerlengo, Joshi and Kumar ^33^ investigated the oral microbial profile of 192 participants belonging to Chinese, Latinos, non-Hispanic whites, and non-Hispanic blacks, finding the lower alpha diversity of non-Hispanic blacks than the other three groups. Yang, Zheng, Cai, Shrubsole, Pei, Brucker, Steinwandel, Bordenstein, Li and Blot ^34^ observed the differences in overall microbial composition between African Americans and European Americans. A recent study documented different microbial alpha and beta diversity of supragingival plaque microbiome among 96 children from four ethnic groups, including African American, Burmese, Caucasian and Hispanic ^35^. A study conducted by our group demonstrated the differences between Australian and Chinese wine experts’ salivary protein profile and their sensory responses ^36^. The mode of variation between Australian and Chinese saliva has not been established, but our preliminary data suggest that dietary differences might be correlated with these differences (data not shown).

Food preference is strongly influenced by a consumer’s cultural, geographic and personal background ^37, 38^. The preferences of Australian and Chinese consumers for 14 red wines have been compared by Williamson, Robichaud and Francis ^37^, finding variance in some attributes for liking, such as “alcohol”, “sweetness” and “dark fruit”. Aroma perception is a prominent factor that influences the acceptability and preference of customers for different wines ^39^. However, much uncertainty exists about the relationships between host factors, salivary composition, and aroma. We ask if the functional properties of diverse salivary microbiome communities can provide insight into the differences in sensory perception amongst wine consumers.

To enhance our understanding of saliva composition and aroma perception, we analysed samples from 30 participants from Australia and China and mixed the saliva samples with wine to observe the ability of human saliva to release aroma compounds from wine. Each participant provided three types of saliva samples collected before, during and after mechanical stimulation to extend our knowledge of reactions that occur in the oral cavity during stages of appreciating wine. Enzymatic activity tests and full-length 16S rRNA sequencing highlighted the differences of saliva type and origin of participants on salivary composition. Combining these findings with the wine aroma profile detected by headspace solid-phase microextraction gas chromatography-mass spectrometry (HS-SPME-GC-MS), we demonstrate differences in salivary proteins and microbiome due to the mode of salivary stimulation and how this varies based on geographic origin of the participant.

## 2 Material and Methods

### 2.1 Materials and reagents

All reagents and chemicals used in this study were of analytical grade. Methanol, citric acid, 4-Nitrophenyl acetate, sodium chloride, sodium phosphate, sodium bicarbonate, potassium chloride, potassium phosphate dibasic trihydrate, calcium chloride dihydrate, mucin from porcine stomach, phosphate-buffered saline, Tris-buffered saline, 4-octanol, amylase activity assay kit, and standards for volatile compounds detected in GC-MS analysis was obtained from Sigma-Aldrich (Castle Hill NSW, Australia). Micro bicinchoninic acid (BCA) protein assay kit was acquired from Thermo Fisher Scientific (Waltham, MA). Pure water was obtained from a Milli-Q system (Millipore Australia, Bayswater, Victoria, Australia). A Shiraz wine (Ladies Who Shoot Their Lunch; vintage 2019; 14.9% v/v alcohol) produced in Victoria, Australia, was purchased from a local wine producer, and used in all GC-MS assays with wine (section 2.6). This wine was selected because different aroma perception were observed between Chinese and Australian panellists in a previous sensory evaluation ^36^.

### 2.2 Study cohort

We recruited 30 healthy adults to meet the following criteria: (i) self-reported good systemic health (i.e., no diagnosed diabetes, cardiovascular diseases, endocrine disorders, or hypertension); (ii) no clinically diagnosed oral diseases; (iii) not receiving antibiotic or steroid hormone use within the previous three months; (iv) no smoking habits (less than several times a month); (v) not a regular drinker (less than several times a week); (vi) have a BMI index no higher than 24 kg/m^2^. The sample collection was performed at the University of Melbourne, Australia. An online questionnaire was provided to all volunteers to ensure they met the inclusion criteria. Demographics were also acquired using self-reported questionnaires before the collection. Fifteen Chinese and fifteen Australian participants (aged 18-28) joined the study. All participants resided in Melbourne, Australia. The participants were grouped according to their country of birth and residence history. The Australian group includes participants born in Australia and have spent most of their lives there. The Chinese group contains participants born in China and have lived in Australia for less than five years. The ethnic origin of the participant’s parents or grandparents was not collected nor considered in this study. Participants self-identified as ‘Chinese’ or ‘Australian’. The gender and BMI index of participants were balanced in each group **(Table S1)**. This study was approved by the University of Melbourne Human Research Ethics Committees (Review Reference Number: 2022-24220-29889-3).

### 2.3 Sample collection

Before collection, sampling procedures were explained in detail to all participants. Participants were asked to refrain from eating and drinking any beverage except water for an hour prior to providing saliva samples. Three types of saliva were collected before (Phase A), during (Phase B), and after (Phase C) stimulation. The collection procedure is illustrated in **Figure S1**. For Phase A, participants naturally expectorated the unstimulated saliva into a 50 mL sterile collection tube until the volume reached 5 mL. For Phase B, participants were instructed to chew a piece of paraffin wax and expectorated the stimulated saliva into another collection tube in 5 minutes. Finally, after removing paraffin wax from the oral cavity, the participant repeated the saliva collection procedure described in Phase A. The saliva samples collected after stimulation were named as Phase C. Time was recorded in Phases A and C to calculate the flow rate as volume collected per minute (mL/min). For the same participant, Phase A, B, and C collections must be completed consecutively on the same day. Immediately after the collection, saliva samples were separated in smaller aliquots and kept frozen at -80 °C until analysis.

### 2.4 Biochemical analyses

Salivary total protein content, and enzymatic activities of 90 samples were tested individually to find the differences among different types of saliva. All analyses were performed in triplicates. Protein concentration of whole saliva samples was tested by Micro BCA™ Protein Assay Kit (Thermo Scientific™) using bovine serum albumin as standard for calibration.

For the enzymatic activity measurements, saliva samples were centrifuged at 15,000 ×g for 15 min at 4 °C to remove insoluble matter. All enzymatic activity assays were performed in triplicate. Total salivary esterase activity was conducted on individual samples accordng to the method from Pérez-Jiménez, Muñoz-González and Pozo-Bayón ^26^. α-Amylase activity assay was performed following the user manual of an Amylase Activity Assay Kit (Sigma-Aldrich), we pooled the samples for this assay because of the limited material for some participants.

### 2.5 Full length 16S rRNA gene sequencing

Bacterial genomic DNA was extracted from 90 whole human saliva samples using the QIAGEN® DNeasy® PowerSoil® Kit. The concentration of genomic DNA was measured using a Qubit HS DNA kit (Thermo Scientific™) and normalised to 1 ng/μL prior to Polymerase Chain Reaction (PCR). Following PacBio’s recommended PCR protocol, full-length 16S amplicons were generated using the Platinum^TM^ SuperFi II High-fidelity PCR enzyme (Thermo Scientific™), with 2ng of input DNA. The amplicons generated were visualised on a 2% agarose gel and pooled. SMRTbell^TM^ cleanup beads (PacBio) were used to purify the pooled amplicons according to the PacBio protocol. The amplicon library was contracted with the SMRTbell® prep kit 3.0 library prep kit (PacBio) following the protocol. The library was then polymerase-bounded using the Revio^TM^ Polymerase Kit (PacBio) and loaded on a single-molecule real-time (SMRT) Cell 25M (PacBio). 16S rRNA full-length sequencing was performed for 30 hours on the PacBio Revio instrument.

### 2.6 Gas chromatography-mass spectrometry (GC-MS) analysis for whole human saliva

Different types of saliva samples were pooled together for the headspace solid-phase microextraction GC-MS analyses (HS-SPEM-GC-MS) to overcome the inter-individual variation in saliva composition and the availability of saliva from each participant. Saliva was pooled based on stimulation phase, by participant origin, resulting in 6 groups of saliva samples: Australia-Phase A, Australia-Phase B, Australia-Phase C, Chinese-Phase A, Chinese-Phase B and Chinese-Phase C. Each sample type was measured in triplicate.

Three types of controls were used: the mixture of wine and water, the mixture of wine and artificial saliva, and the mixture of six pooled saliva samples and model wine. The mixture of wine and water were used to control for the dilution effect of saliva in the GC analysis matrix ^4, 40^. An artificial saliva solution was spiked with wine to control for the effect of mucin and salts in aroma release. It was prepared according to the method of Genovese, Piombino, Gambuti and Moio ^1^. The artificial saliva samples were freshly prepared by dissolving mucin (2.16 g/L), NaHCO_3_ (5.208g/L), K_2_HPO_4_·3H_2_O (1.369 g/L), NaCl (0.877 g/L), KCl (0.477 g/L) and CaCl_2_·2H_2_O (0.441 g/L) in Milli-Q water. The mixture of the model wine and six types of saliva were used to control for the confounder effect of the volatiles that may present in saliva itself. The model wine used in this study was prepared by saturating 12% ethanol (v/v) with potassium hydrogen tartrate, adjusting to pH 3.0 by 40% (w/v) tartaric acid. Each mixture was measured in triplicate.

The static headspace volatile extraction method was conducted following the method of Muñoz-González, Feron, Guichard, Rodríguez-Bencomo, Martín-Álvarez, Moreno-Arribas and Pozo-Bayón ^41^ with some modifications. The 20 mL GC vial was incubated at 37°C for 20 min to stabilise the temperature of the reaction environment. Then 5 mL of wine and 1 mL of pooled saliva were added to the vial with 10 μL of internal standard (100 mg/L 4-octanol). The wine-saliva mixture was incubated at 37 °C for 12 min. The headspace volatile extraction from the GC vial was performed by a 65 μL polydimethylsiloxane/divinylbenzene (PDMS/DVB) SPME fibre with 1 cm fibre length (Supelco, Bellefonte, PA) at 37 °C for 5 min.

GC-MS analysis was carried out with an Agilent 7890A GC system (Agilent Technologies, Mulgrave, VIC, Australia) coupled with an Agilent 5975C MS (Agilent Technologies). The fibre was desorbed at 220 °C in splitless mode at 8.7 psi for 30 s. Chromatographic separation of volatile compounds were performed with a DB-Wax polar capillary column (Agilent Technologies; 30 m × 0.25 mm × 0.25 μm film thickness). Helium was used as the carrier gas at a flow rate of 0.7 mL/min. After initially holding the oven temperature at 40°C for 4 mins, it then ramped at 3°C/min to 220 °C, and held at 220 °C for 10 min. For the MS system, electron ionisation mode at 70 eV was used to record the electron impact mass spectra. The scanning mass range was from m/z 35 to 350. The temperature of the MS interface, MS source, MS quadrupole was 250, 230 and 150 °C.

The volatile compounds identification was based on the retention indices (RIs) with the facility of NIST library (version 11.0). RIs were obtained by calculating the retention times of alkane standards (C_5_–C_30_) on the same column. Peaks were semi-quantified using the internal standard, 4-octanol. The concentrations of volatiles were expressed as μg/L 4-octanol equivalence. Among the 27 compounds identified, sixteen compounds have also been quantified using external standards to validate the results of the semi-quantified method, facilitated by the calibration curve of standard compounds as:

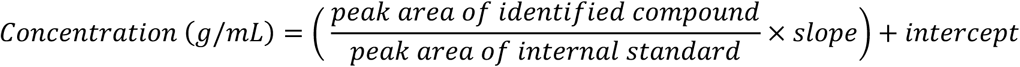

### 2.7 Statistical analyses

The data analyses of salivary biochemical parameters and the volatile compounds generated from the wine-saliva mixture were conducted in R (version 4.1.0). Student t-tests were used to examinate the significant (*p* < 0.05) differences between samples from Chinese and Australian participants. Two-way analysis of variance (ANOVA) with Tukey’s comparison were performed to test the significant (*p* < 0.05) differences among saliva samples collected before, during and after stimulation. The Spearman’s correlations were calculated with *rcorr* function in the R package *Hmisc* ^42^.

For bioinformatic analyses, the demultiplexed PacBio HiFi full-length 16S rRNA gene sequences are quality filtered using QIIME2 ^43^. DADA2 ^44^ was used to denoise the sequences into amplicon single variants (ASVs) using the function “dada2 denoise-ccs”. A phylogenetic tree was generated from the *mafft* alignment. The naïve Bayesian machine learning based classification was performed using the expanded Human Oral Microbiome Database (eHOMD) version 15.23 ^45^. If a species-level match was not found, it was further classified based on three other databases that successively refer to the next one. The three databases are GTDB r207, Silva v138 ^46^ and the NCBI RefSeq 16S rRNA database supplemented by the Ribosomal Database Project (RDP). The resulting objects were imported to R (version 4.1.0) for further analyses. Alpha diversity was calculated with Simpson’s diversity index.

Wilcoxon rank-sum tests were used to determine the statistically significant (*p* < 0.05) differences. Beta diversity was assessed using Bray-Curtis dissimilarly. Permutational multivariate analysis of variance (PERMANOVA) with 999 permutations was conducted to examine the statistically significant (*p* < 0.05) difference in beta-diversity using the *adonis2* function of *vegan* package ^47^. The differential taxa were defined from phylum level to species level by *metacoder* package ^48^ with false discovery rate-adjusted Wilcoxon rank-sum test. ANCOM-BC2 ^49^ was performed to identify the differential species with the default parameters.

## 3 Results

### 3.1 Simulated saliva (Phase B) has a distinguishable protein and biochemical composition

#### 3.1.1 Differences in saliva composition amongst the three phases

To determine whether human saliva produced before, during and after stimulation is different, the biochemical properties and microbial composition of human saliva samples were tested. The salivary flow rates were calculated as volume collected per minute, reflecting the speed of participants to produce saliva. *Ex vivo* measurements of α-amylase and esterase were conducted to assess enzymatic activity of the centrifuged saliva. The flow rate observed in Phase B and total salivary esterase activity were significantly higher than the other two phases, while Phase A and C saliva were not significantly different **(Figure 1A, E)**. For the total protein concentration of saliva samples, the value of Phase A was significantly higher than that of the other two phases **(Figure 1C)**. No significant differences were found among the three groups in the amylase activity **(Table S2)**.

**Figure 1.** The differences of salivary composition amongst three phases of collection and two participant groups. Boxplots showing the flow rate of (**A**) Phase A, B, C and (**B**) Australian and Chinese participants; the total protein concentration of (**C**) Phase A, B, C and (**D**) Australian and Chinese participants; the total salivary esterase activity of (**E**) Phase A, B, C and (**F**) Australian and Chinese participants. Significant differences (*p*< 0.05, Student’s t-test; *p*< 0.05, one-way ANOVA) were tested and are marked with “*”; with NS denoting no significant differences observed.

Full-length 16S rRNA gene sequencing was used to detect the differences in the salivary microbiome. In contrast to the protein concentration and enzymatic activities, no significant differences were detected among Phases A, B and C regarding the alpha-diversity and beta-diversity of the microbial communities (*p* > 0.05). Although several species have been identified as differential taxa **(Figure S2)**. Most differential species had the lowest relative abundance in Phase B **(Figure S2A)**. The differential species identified in Phase A and C were clustered together in the heatmap **(Figure S2B)**, suggesting the microbial profile is distinct in Phase B at species level.

As the volatiles may also have originated from the human oral cavity, the volatiles in the saliva samples were assessed by mixing each pooled saliva sample with model wine. Among the detected volatiles, (–) – clovene was found in the mixture of saliva and wine and the mixture of saliva and model wine. Therefore, this compound was excluded from further analysis.

#### 3.1.2 Effect of stimulation on the release of aroma from wine

To evaluate the ability of different saliva samples to release aroma compounds from wine, HS-SPME-GC-MS analyses were performed on the same wine sample spiked with saliva samples. Saliva samples were pooled into six groups: Australia-Phase A, Australia-Phase B, Australia-Phase C, Chinese-Phase A, Chinese-Phase B and Chinese-Phase C. After mixing the six pooled saliva samples with wine, twenty-seven volatile compounds were identified, consisting of 12 esters, 6 alcohols, 3 acids, 2 benzenoids and 4 terpenes **(Table 1)**. The wine sample was also mixed with water and artificial saliva as the control groups **(Figure S3A)**. For many compounds, especially esters, the addition of artificial saliva induced a significant decrease in their concentration compared to the wine-water mixture. Depending on the type of pooled saliva added, the decrease caused by artificial saliva could either be counteracted or further decreased **(Figure S3B)**. It suggests the whole human saliva induces other reactions with the wine matrix other than the effect of mucin.

**Table 1.**
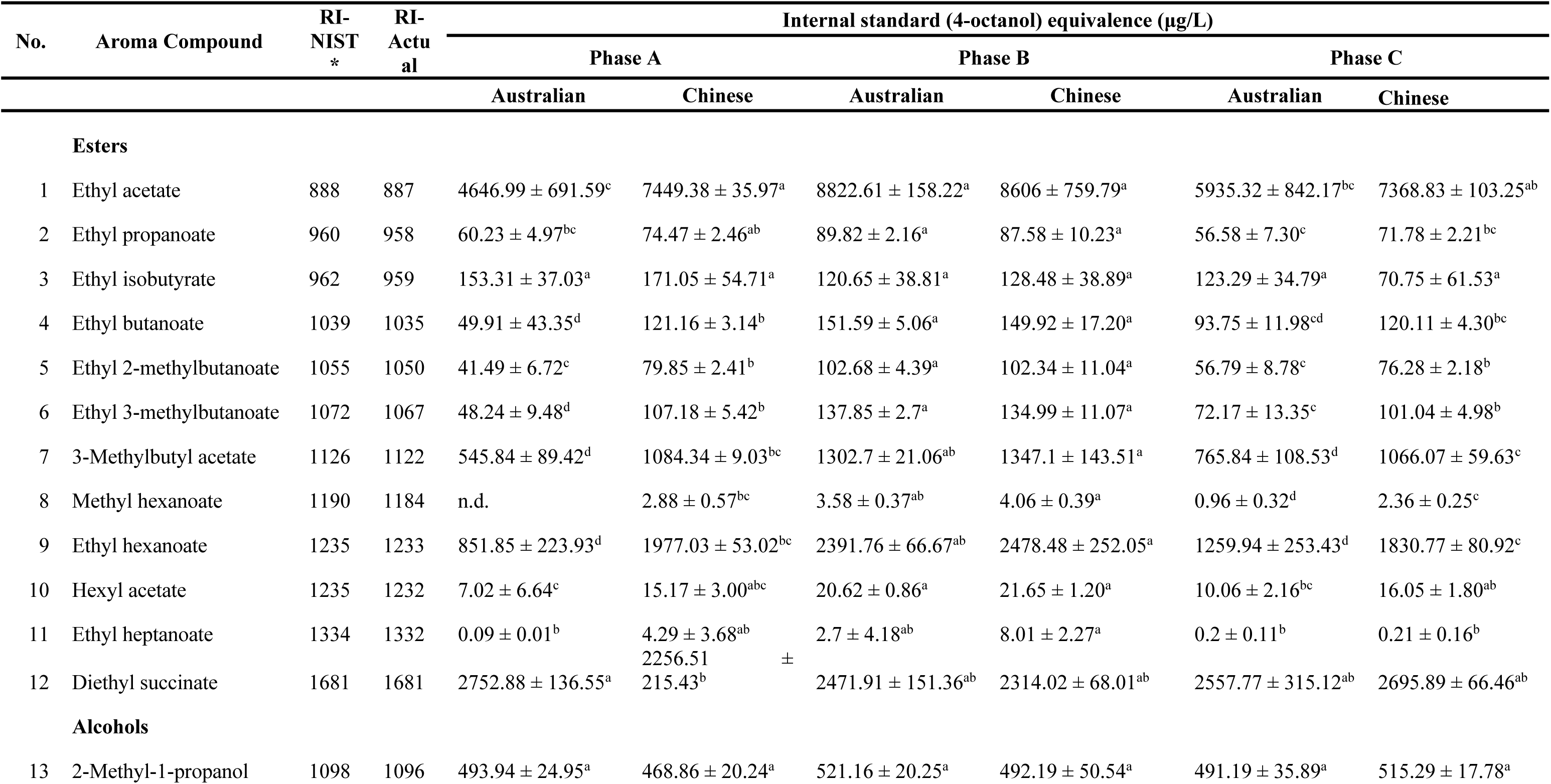
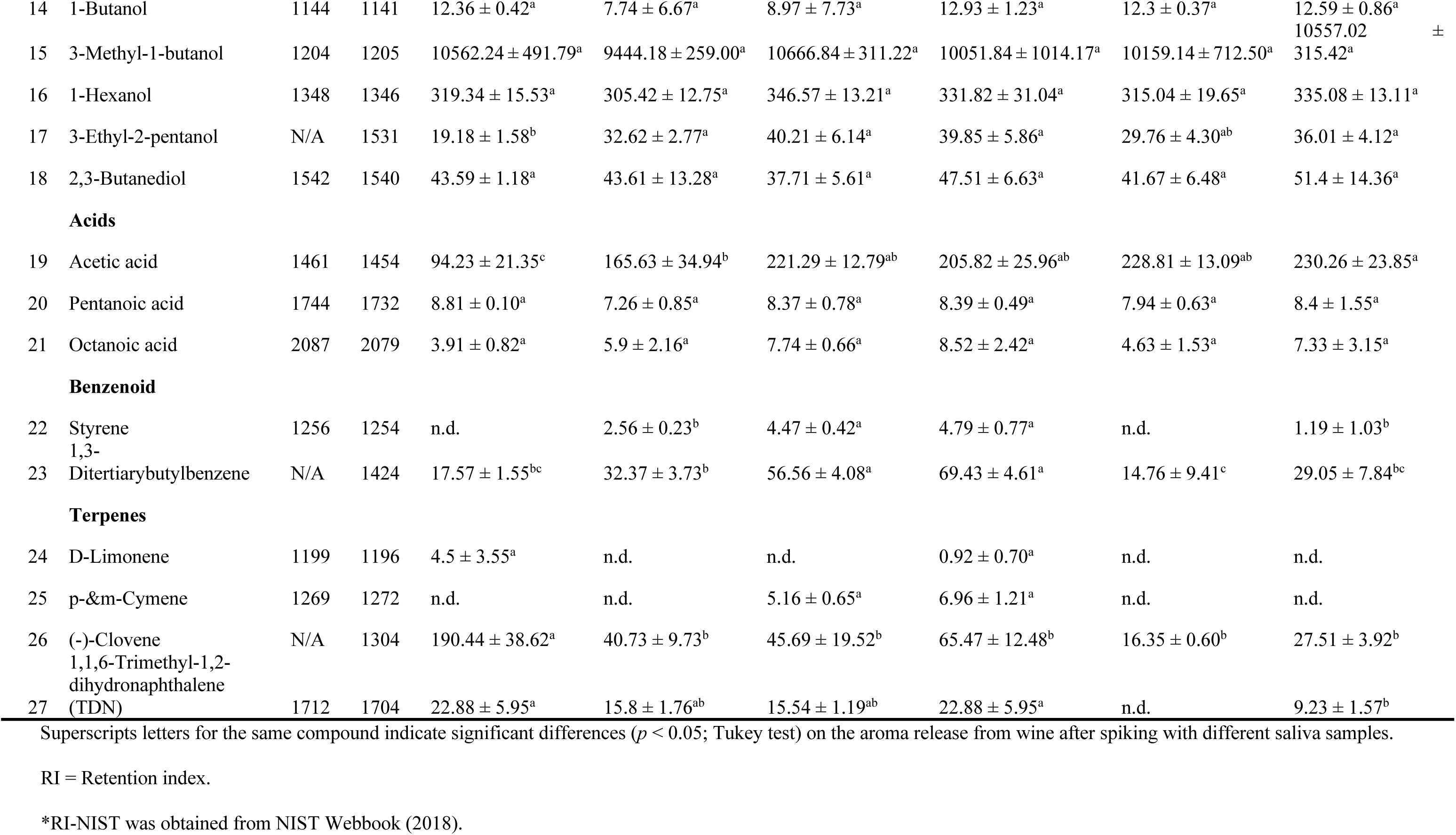
Aroma compounds detected by headspace solid-phase microextraction gas chromatography mass spectrometry (HS-SPME-GC-MS) from the mixture of six types of pooled saliva samples and wine.

For the 26 included volatiles, twenty-one volatiles were constantly detected across all six groups, while the other 5 volatiles were only occasionally found **(Table 1)**. Methyl hexanoate is the only ester that was not constantly detected in all groups. The other occasionally detected compounds are all benzenoids and terpenes. These compounds were mostly absent in Phases A and C samples except limonene, which was not detected in the Australian-Phase B group. The one-way ANOVA conducted with the concentration of 16 volatiles quantified using external standards is compatible with the results calculated using semi-quantification method **(Table 1; Table S3)**.

The dataset was divided into two subgroups, Australian **(Figure 2A)** and Chinese **(Figure 2B)**, to show the changing trend for 21 constant-detected compounds among Phases A, B and C. The statistical analyses suggest that the Australian and Chinese groups individually have 9 and 8 compounds with significantly higher concentrations in Phase B than in the other two phases (*p* < 0.05). Most of the distinguishing compounds are esters, while octanoic acid and 1,3-ditertiary butyl benzene show the same trend in either the Australian or both groups. In addition, several compounds demonstrated significant differences between Phase A and C, including diethyl succinate, 3-methylbutyl acetate, 3-ethyl-2-pentanol and acetic acid. 3-Ethyl-2-pentanol and acetic acid only showed a significant difference among phases in the Australian group, with significantly lower concentration in Phase A (*p* < 0.05). Further, a significant difference was observed between the Australian and Chinese groups of these two compounds in Phase A **(Figure 2C)**.

**Figure 2.** The aroma released from wine with saliva samples from three collection phases are different. Point plots with lines showing the change of each volatile released from (**A**) wine-Australia saliva mixture and (**B**) wine-Chinese saliva mixture. The green lines represent the compounds showing highest concentration in Phase B and have no significant differences between Phase A and C (one-way ANOVA, *p* < 0.05). Among them, light green lines represent compounds only significantly higher than Phase A, while dark green lines represent compounds significantly higher than both Phase A and C. The pink lines represent the compounds that also demonstrate significant differences between Phase A and C. The dark pink lines showing the compounds have significant differences among all three comparisons. The light pink lines showing the compounds have differential concentration between Phase A and B, and Phase A and C. The volatiles with significantly changes are indicate by letters: a) Ethyl acetate; b) Diethyl succinate; c) Ethyl hexanoate; d) 3-Methylbutyl acetate; e) Ethyl butanoate; f) Ethyl 3-methylbutanoate; g) Ethyl propanoate; h) Ethyl 2-methylbutanoate; i) Hexyl acetate; j) 3-Ethyl-2-pentanol; k) Acetic acid; l) 1,3-Ditertiarybutylbenzene; m) Octanoic acid. (**C**) Bar plots showing the concentration of 3-Ethyl-2-pentanol and Acetic acid in six subgroups. Significant differences (*p*< 0.05, Student’s t-test) between Australian and Chinese group in Phase A are marked with “*”. (**D**) Principal component analysis (PCA) plot demonstrating the wine aroma release after spiking with samples from three groups.

To better understand the relationship between three types of saliva and the aroma released from wine, a principal component analysis (PCA) was performed on the 21 detected compounds shared amongst all samples **(Figure 2D)**. The internal standard equivalent concentration of the aroma compounds, (expressed as μg/L 4-octanol equivalents), distinguished the saliva samples from different phases. The first two principal components explained 68.57% of the variance. The first principal component (PC1) separated the samples of Phase B from those of Phase A and C. The samples of Phase B were highly associated with most esters, 1,3−ditertiarybutylbenzene and 3−ethyl−2−pentanol, positioned in the space defined by negative PC1. On the contrary, 1-butanol was positively related to PC1. The second principal component (PC2) was negatively associated with most alcohols, including 2-methyl-1-propanol, 3-methyl-1-butanol, 1-hexanol and 2,3-butanediol. Phase A and C samples were separated on PC2 with some overlap retained, suggesting the interaction between the two investigated covariates, phase, and subject origin **(Table S2)**. In addition, PCAs were conducted using a total 27 compounds and 16 compounds quantified using external standards, separately **(Figure S4)**. These results agree with the PCA constructed by the semi-quantified compounds.

### 3.2 The composition of saliva from Australian and Chinese participants is different

#### 3.2.1 Differences in salivary proteins between two groups

The flow rate, total protein concentration, and activities of alpha-amylase and esterase of the saliva samples collected from Australian and Chinese participants were compared to investigate the influence of the host’s cultural background on salivary composition. Although no differences were found between the flow rate of Australian and Chinese participants’ saliva **(Figure 1B)**, the saliva of Australian participants demonstrated higher total protein concentration **(Figure 1D)**. For the enzymatic activities, the two groups demonstrated no statistical differences in either amylase or esterase activity **(Table S2; Figure 1F)**.

#### 3.2.2 Differences in salivary microbiome profiles

For the comparison of the salivary microbial profile of participants from Australia and China, the full-length 16rRNA gene was sequenced. After denoising by DADA2, the samples were rarefied to 5,232, resulting in 11,052 amplicon sequence variants (ASVs). The Australian and Chinese groups differ in the alpha-diversity characterised by Simpson Index **(Figure 4A)**. Beta-diversity analysis conducted using Bray-Curtis dissimilarity also demonstrated the significant differences between these two groups (PERMANOVA, *p* = 0.001). The principal coordinate analysis (PCoA) result showed the separation between the microbial composition of samples from two groups **(Figure 4B)**.

When investigating the taxa that discriminate saliva samples from Australian and Chinese participants, we first compared the relative abundance of the two groups at phyla **(Figure 3A)** and genus levels **(Figure 3B)**. Firmicutes, the phylum with the highest relative abundance in both groups, showed higher relative abundance in the saliva samples from Australian participants. A higher relative abundance of *Veillonella* in Australian saliva was observed. We performed a Wilcoxon Rank Sum test with a false discovery rate correction to each taxon, spanning all taxonomic levels **(Figure 3C)**. Genus *Neisseria*, *Lautropia*, *Porphyromonas*, *Capnocytophaga*, *Cardiobacterium*, *Filifactor* and *Parvimonas* were higher in Chinese participants’ saliva, with several species of these genre. All taxa belong to phyla Synergistetes and Spirochaetes, except the species were more prevalent in the Chinese group. In contrast, the saliva microbiota of Australian participants was distinguished mainly by taxa under the order *Veillonellales*, especially the genus *Veillonella* and *Megasphasera*. The genus with high abundances, such as *Streptococcus*, *Prevotella* and *Haemonphilus*, contain both species identified as biomarkers of Chinese and Australian participants.

**Figure 3.** The microbial profiles of Australian and Chinese participants’ saliva samples are different. Comparison of the salivary microbial composition between two groups in stacked bar plots at (**A**) phylum level and (**B**) genus level. (**C**) Taxonomic hierarchies showing the enrichment of taxa from two groups from phylum to species level. The taxa with significant differences (false discovery rate-adjusted Wilcoxon rank-sum *q* < 0.05) were coloured with green (higher in Australian group) and pink (higher in Chinese group).

**Figure 4.** The differences in salivary microbial composition between Australian and Chinese participants associate with the differences in aroma released after mixing with wine. (**A**) Comparison of the alpha-diversity between two groups, representing by Simpson’s Diversity Index (Wilcoxon rank-sum test; *p* = 0.029). (**B**) Principal coordinate analysis (PCoA) with Bray-Curtis dissimilarity showing the difference between two groups (PERMANOVA; *p* = 0.001). (**C**) and (**D**) Heatmap represents the mean relative abundance of (**C**) eight differential species identified by ANCOM-BC2 and (**D**) three species that significantly correlated (p < 0.01, FDR-adjusted Spearman’s rank correlation) with aroma compounds in six pooled saliva groups. (**E**) Bar plots showing the log-transformed concentration (μg/L 4-octanol equivalence) of aroma compounds detected in two groups. The coloured squares on the bottom indicate the result of two-way ANOVA. (**F**) Spearman’s correlation heatmap between aroma compounds and relative abundance of species. Significant correlated pairs (*p* < 0.01, FDR-adjusted Spearman’s rank correlation) are labelled with “*”.

We also used ANCOM-BC2 to identify the differential species between Australian and Chinese saliva to accomplish more precise results at the species level. We found 8 differential species for comparing two groups, mostly under the phylum *Bateroidetes* and *Firmicutes* **(Figure 4C)**. Among them, *Porphyromonas pasteri* and *Neisseria subflava* have higher relative abundance in the Chinese group, which agrees with the results of the Wilcoxon Rank Sum tests **(Figure 3C)**. Similarly, the six differential species enriched in Australian samples identified by ANCOM-BC2 are also found in the Wilcoxon Rank Sum tests. In the Australian group, the differential species of Phase A and C were clustered together, suggesting they are closer to each other than Phase B. Meanwhile, the differential species in the Chinese Phase A and B samples were clustered.

#### 3.2.3 Differences in saliva composition between groups are associated with aroma release

To investigate the aroma consequences of compositional differences in the saliva of saliva samples collected from Australian and Chinese participants, the volatile profile was collected by HS-SPME-GC-MS and compared. The result of two-way ANOVA showed that 11 compounds were released differently after mixing with saliva samples from two groups. Different concentrations due to the saliva samples collected at different phases were observed **(Figure 4E)**. All differential compounds showed higher concentration in the Chinese group, especially in unstimulated Phases A and C. Meanwhile, ethyl heptanoate released after mixing with Chinese saliva samples demonstrated higher concentration in Phases A and B.

We suspected that the salivary bacteria contributed to aroma release, and so we calculated the Spearman’s rank correlation between the relative abundance of the species in pooled saliva samples and the aroma compounds released after mixing the wine samples with those saliva samples. We found three species significantly correlated to wine aroma profile, including *Streptococcus sp. HMT006*, *Prevotella shahii* and an unclassified species belongs to *Selenomonas* **(Figure 4D)**. The concentration of three differential esters, ethyl propanoate, 3-methylbutyl acetate and ethyl hexanoate, are positively associated with the relative abundance of *Streptococcus sp. HMT006*, while negatively associated with the relative abundance of the unclassified species of *Selenomonas* **(Figure 4F)**. The higher concentration of 3-ethyl-2-pentanol and several esters in the Chinese group has been positively correlated with the increased relative abundance of *Streptococcus sp. HMT006* in saliva samples from Chinese participants in Phase A. As the only benzenoid identified, the concentration of 1,3-ditertiarybutylbenzene showed a negative correlation with the unclassified species of *Selenomonas.* Although two-way ANOVA concluded that the concentration of 2,3-butanediol was not different between the Chinese and Australian groups, it was positively associated with the higher relative abundance of *Prevotella shahii* in the Chinese saliva

## 4 Discussion

In the current study, we observed the differences in salivary composition between saliva samples collect at different stimulation stages and from participants with different geographic origin. This study further demonstrates the association between host factors, salivary composition, and aroma release from wine. For the two parameters controlled in this study, the phase of saliva production and the host’s ethnic background, we simultaneously observed the differences in salivary composition and the aroma profiles of wine after mixing with saliva samples. The salivary stimulation stages and subject origin both influence aroma release ability as the composition of saliva is different. These findings shed light on the role of human saliva in immediate and prolonged retronasal perception of aroma and has introduced unstimulated saliva as a control in *ex vivo* experiments.

Overall, saliva samples collected before, during and after stimulation were different in composition. We observed a higher total protein concentration in the saliva samples produced in Phase A. The lower protein concentration in Phase B and C may be due to the dilution of the substantial water content secreted in Phase B ^50^. Interestingly, even though the flow rate dropped significantly after removing the stimulus from the oral cavity, the total protein concentration in saliva did not return to the level when the stimulus was not received. Further analyses, such as proteomics should be applied to assess the decreasing rate of individual proteins. By observing whether the decreasing rate of different proteins are even, it is possible to infer if dilution is the main reason for the lower protein concentration in Phase C. In contrast, a higher total salivary esterase activity was found in Phase B, while the esterase activity of Phases A and C had no significant difference. The esterase in saliva is an important enzyme for hydrolysing esters into carboxylic acids and alcohols ^6^. The higher esterase activity in stimulated saliva than in unstimulated saliva has also been observed by Pérez-Jiménez, Muñoz-González and Pozo-Bayón ^26^. Esterase did not follow a consistent trend with total protein concentration, suggesting the mechanisms beyond dilution that may be responsible for modulating the concentration of different proteins after stimulation. Meanwhile, no significant differences were observed in the current study between the microbial diversities of saliva samples collected at different stimulation stages. The association between oral microbiota and the type of saliva has been widely investigated by previous studies but remains controversial. A previous study suggested a significant difference between unstimulated and stimulated saliva in both alpha and beta diversity ^22^. On the contrary, another two studies ^23, 24^ reported no significant differences between the Shannon diversity index of bacteria in unstimulated and stimulated saliva samples. Several species have been identified as differential among the mouth-rinse water, unstimulated saliva, and stimulated saliva by Jo, Nishimoto, Umezawa, Yama, Aita, Ichiba, Murakami, Kakizawa, Kumagai, Yamada and Fukuda ^24^, which agrees with our results. However, no intersection has been found between the differential species defined by the previous work and our study. The reason could be the different statistical tools used. It is possible that the differences could be too subtle to detect. Further study with higher resolution, such as metagenomic sequencing, need to be applied to monitor the sub-species level or community-level differences.

The relationship between the stimulation stages and aroma release ability was found to be focused on the esters in wine and the esterase activity in human saliva. Esters are an essential group of aromatic compounds to contribute fruitiness to wine ^55^. Most esters detected in this study have the odour activity values greater than 1 **(Table S3)**, meaning that these compounds are likely to have a sensory impact. Saliva samples produced in Phase B demonstrated the highest esterase activity and induced the highest concentration of esters released from wine samples. However, the presence of esterase was previously associated with the hydrolysis of esters ^3^ and the generation of carboxylic acids ^11^. Esters are relatively less persistent in the oral cavity after swallowing, probably due to the modification by oral microbiota directly or with enzymatic action ^56^. This rather contradictory result may be explained by the presence of reactions other than enzymatic hydrolysis of the esters. In addition, the total salivary esterase activity could be attributed to different types of enzymes, such as carboxylesterases, carbonic anhydrases, trypsin, and cholesterol esterases ^26^. The catalytic effect of specific enzymes could be accompanied by concurrent synthesis of esters. Interestingly, the same phenomenon was reported in our recent study ^36^, which strengthens our assumption that perhaps more than one type of reaction involving esterases occurs when human saliva mixes with esters in wine.

Notably, several volatiles showed differences between Phase A and Phase C in the Australia group, including 3-methylbutyl acetate, 3-ethyl-2-pentanol and acetic acid. 3-Methylbutyl acetate is an ester with fruity, banana-like aroma (**Figure S3**). In contrast, 3-ethyl-2-pentanol and acetic acid may provide an unpleasant flavour to wine ^57^. Those volatiles may play a role in the after-stimulation stage. During retronasal perception of wine, two distinct types of aroma release are perceived by consumers: the immediate impression produced when swallowing and the prolonged perception after swallowing ^58^. By comparing the saliva samples collected during and after stimulation with the control, our study demonstrates that it is important to specify the type of saliva collected and used in *ex vivo* experiments. Stimulated saliva needs to be specified for the research targeting the reactions with stimulus. These findings could also shed light on understanding the differences between immediate and prolonged perception of wine aroma. Meanwhile, although we demonstrated the different aroma release capacities of saliva samples collected during and after the mechanical stimulation, more complex mechanisms are introduced in aroma perception occurs during the wine tasting experience of human beings. Mixing the saliva samples collected after the mechanical stimulation with wine could not completely reflect the complexity and dynamics of “after-odour” occurring in human oral cavity. We set the stopping point for the collection of Phase C as the time that volume reaches 5 mL to allow enough time for saliva to change from “stimulated” to “unstimulated”. However, after swallowing the wine, various amounts of odorants will be captured by oral mucosa and keep working as “stimulus” in mouth ^28^. Therefore, the changes of saliva from “stimulated” to “unstimulated” could not be as abruptly as what we mimicked using paraffin chewing. The proportion of wine and saliva used in our static headspace SPME method does not reflect all the consequences of oral processing to explain aroma production in the ‘after-taste’ period. Further analytic approaches to simulate the relationships between post-stimulated saliva and wine, include modelling the oral mucosa ^60^, and should be used to further understand the role of saliva in the after-taste phase.

Moreover, the chemical composition of wine also influences the aroma release of wine aroma volatiles. For example, the phenolic compounds impact the aroma release depending on the chemical structure ^61,62^. Specifically, phenolic acids facilitate the intra-oral release of β-phenyl ethanol, linalool and β-ionone, while flavonoids in wine decrease the release of some esters ^61^. Meanwhile, ethanol in wine may also affect the salivary proteins ^63^. Future research is required to explore the influences of host factors on the aroma release ability of saliva on other food matrices, such as non-alcoholic beverages and solid foods.

The influence of the host’s origin on salivary composition, especially the oral microbiome, has also been observed in the current study. To control the confounder effect of other host factors, we applied a series of selection criteria to ensure the two groups of participants have balanced gender and BMI. Although significant difference was identified in age of two groups, all participants in this study were aged between 18 to 28. The differences induced by host origin have been attributed to genetic or external factors, such as diet ^51^ and altitude ^52^. Differences in cultural background and lifestyle habits between Asian countries and Western countries have led to interest in understanding the differences in salivary composition between different countries ^53^. Our previous study collected salivary microbiome data from 47 publicly available datasets, reporting the differences in beta diversity of saliva samples from three geographic locations, including North America, Europe, and China ^16^. Notably, the variance explained in the previous study could be confounded by external factors other than the differences among three populations, such as climate and collection methods. To better understand the differences between two participant groups, the current study effectively eliminated the variance caused by geographic location by collecting all the saliva samples from the same location within a short period. *Veillonella* spp., including *Veillonella atypica* and *Veillonella dispar*, were identified as the differential taxa that were higher in Australian samples, which was consistent with the result of our previous study. Among them, *Veillonella atypica* has also been identified as a biomarker to differentiate people with obesity from normal-weight individuals ^54^, underlying a potential association among diet, body weight and ethnicity.

Although the differences between the Chinese and Australia group have been observed in our study, association between ethnicity and salivary protein composition is far from conclusive. A previous study ^29^ compared protein concentration, flow rate, amylolytic activity, and lipolytic activity of unstimulated saliva samples from 15 Chinese and 15 Caucasian participants living in the Netherlands. Like the current study, they focused on the difference in saliva samples between Chinese and residents of a country defined as “Western countries”. However, they observed higher total salivary protein concentrations in Chinese than in Caucasian participants. This discrepancy between two studies could be attributed to the differences between the Western countries selected. Australian and Dutch residents could have specific differences in diets and lifestyles, leading to the different biochemical parameters of saliva. Meanwhile, no significant differences in flow rate and amylolytic activity was detected by Mosca et al. ^29^ , which is consistent with our findings. Further proteomic analyses on samples collected in the current study would facilitate the observation of the differences on specific protein concentration; we suspect that differences in published studies might also be due to the collection of stimulated or unstimulated saliva (or merging of these two classes).

Our *ex vivo* results show that Australian and Chinese have different aroma compounds released in the oral cavity. Our previous study reported that people from Australia and China have different sensory responses to the same wine ^36^ and reported increased levels of esters in the mixture of Chinese saliva and wine. According to the results of current study, the prevalence of *Streptococcus* sp. HMT 066 has been associated with the concentration of esters released. It has been reported that *Streptococcus* spp. have the ability to release aroma from odourless precursors ^8^. 2,3-butanediol is the second most abundant by-product generated during the alcoholic fermentation of wine, contributing to its bitterness ^64^. The higher 2,3-butanediol concentration found in our study using wine and saliva from the Chinese group positively correlates with the relative abundance of *Prevotella shahii* **(Figure 4F)**. *Prevotella* spp. has been described as a potential contributor to aroma perception by enzymatic activity ^13^. Nevertheless, within genera like *Streptococcus* and *Prevotella*, the function of different species varies. Functional annotation and taxonomic profiling with higher resolution at special or even stain level could be done to differentiate the bacteria belongs to these two genera. Further experiments, such as strain isolation, are required to verify the function of specific species. *Veillonella* spp. have been found different in relative abundance between hosts from China and Western countries in both a previous study ^16^ and the current study. Although no correlation was found between *Veillonella* spp. and any aroma compounds in this study, more research on *Veillonella* could help understand the differences in aroma perception between Chinese and Western people.

In summary, this study demonstrated that saliva secreted before, during and after stimulation differs in composition and the ability of the three types of saliva to release aroma compounds from wine varies. Although Phase A and C have more in common than Phase B, there are still some differences in their composition and the role in aroma release. The enzymatic activity of esterase may explain the variation in the ability of human saliva to release aroma compounds attributed to stimulation. Another important finding of this study was the difference in salivary composition, especially salivary microbiota, between Australian and Chinese participants. Several differential species were identified by full-length 16S rRNA sequencing. We took samples from 30 people in this study, which is a relatively small number, yet our findings mirror data collected in other studies. For example, our findings of a differential relative abundance of *Veillonella* spp. contributes additional evidence to support this genus as a biomarker to differentiate people originating in different geographic areas. The volatiles released from the wine-saliva mixture of these two groups highlight the associations between salivary microbiota and aroma release. Our findings serve as a basis for future studies targeting the relationship between salivary composition, aroma perception, and wine preference. A better understanding of the variations in wine preference gives insights into designing products for specific consumer groups and should be complemented in the future by sensory tests to verify the findings in the human oral cavity.

## Funding

This study was funded by the Faculty of Science at the University of Melbourne. XR and YC are supported by a Melbourne Research Scholarship administered by the University of Melbourne.

## Acknowledgements

The authors would like to acknowledge all participants. The authors would also like to acknowledge the assistance of Zijian Liang to the sample collection process. The work of Dr Pangzhen Zhang to set up methodologies and verify volatile analysis is gratefully acknowledged.

## Ethics approval and consent to participate

The study is under the approval of the University of Melbourne Human Research Ethics Committees (Review Reference Number: 2022-24220-29889-3). All participants provided written informed consent.

## Data availability

The sequencing data generated and analysed in the present study are available in NCBI BioProject database, PRJNA1077481(https://www.ncbi.nlm.nih.gov/bioproject/1077481).

## Declaration of Competing Interest

The authors declare that they have no competing financial interest.

## Supporting Information

**Supplementary Table 1** Background information of participants from Chinese and Australia

**Supplementary Table 2** The influence of two factors, phase and ethnicity on the amylase activity and aroma compounds detected by HS-SPME-GC-MS from the mixture of six types of pooled saliva samples and wine, evaluated by two-way ANOVA and Tukey’s pairwise comparison.

**Supplementary Table 3** The concentration and odour activity values (OAV) of aroma compounds detected by headspace solid-phase microextraction gas chromatography mass spectrometry (HS-SPME-GC-MS) from the mixture of six types of pooled saliva samples and wine, quantified using external standards.

**Supplementary Figure 1** Flow chart showing the sample collection procedure of three types of saliva from each participant: before, during and after stimulation.

**Supplementary Figure 2 Thirteen species identified as differential among three phases by ANCOM-BC. A)** Point plots with lines showing the log-transformed relative abundance of differential species in three phases. The green lines indicate the species with the lowest relative abundance in Phase A, and the pink lines indicate the species with the lowest relative abundance in Phase B; **B)** Heatmap showing the log-transformed mean relative abundance of differential species.

**Supplementary Figure 3 The aroma released from wine after mixing with water, artificial saliva, and pooled saliva samples. A)** The table showing the concentration of volatiles released from wine-water and wine-artificial saliva mixture. Superscripts indicate significant differences (p < 0.05; Student’s t test) between the two groups. **B)** The coloured squares indicate the comparison among wine-water, wine-artificial saliva, and wine-saliva mixtures. The green squares represent the higher concentration of released aroma after mixing with the pooled saliva than mixing with water or artificial saliva. The pink squares represent the lower concentration of released aroma after mixing with the pooled saliva than mixing with water or artificial saliva. The grey squares represent the volatiles that have not been detected. The significant differences (p < 0.05; one-way ANOVA) were indicated by “*”.

**Supplementary Figure 4** Principal component analysis (PCA) plots of A) 27 semi-quantified compounds and B) 17 compounds quantified using external standards showing the similar results with the PCA plots of 21 semi-quantified compounds detected constantly in all subgroups.

## Notes

https://www.ncbi.nlm.nih.gov/bioproject/1077481

### Competing Interest Statement

The authors have declared no competing interest.

### Summary of Updates

Updated after review. Clarifications in the text, extended methods included, figures updated.

