## Supplementary files for "Stimulation of saliva affects the release of aroma in wine; a study of microbiota, biochemistry and participant origin"

**Stimulated saliva has a distinct composition that influences release of volatiles from wine**

The supplementary information contains two parts:

1. **Supplementary Tables**
2. **Supplementary Figures**

**Supplementary Table 1** The influence of two factors, phase and ethnicity on the amylase activity and aroma compounds detected by HS-SPME-GC-MS from the mixture of six types of pooled saliva samples and wine, evaluated by two-way ANOVA and Tukey’s pairwise comparison.

| **No.** | **Aroma Compound or Amylase activity** | ***p* value** | | | | | |
| --- | --- | --- | --- | --- | --- | --- | --- |
|  |  | **Two-way ANOVA** | | | **Tukey’s pairwise comparison** | | |
|  |  | **Phase** | **Ethnicity** | **Interaction** | **A - B** | **A - C** | **B - C** |
|  | **Esters** |  |  |  |  |  |  |
| 1 | Ethyl acetate | p<0.05 | p<0.05 | p<0.05 | p<0.05 | NS | p<0.05 |
| 2 | Ethyl propanoate | p<0.05 | p<0.05 | p<0.05 | p<0.05 | NS | p<0.05 |
| 3 | Ethyl isobutyrate | NS | NS | NS | NS | NS | NS |
| 4 | Ethyl butanoate | p<0.05 | p<0.05 | p<0.05 | p<0.05 | NS | p<0.05 |
| 5 | Ethyl 2-methylbutanoate | p<0.05 | p<0.05 | p<0.05 | p<0.05 | NS | p<0.05 |
| 6 | Ethyl 3-methylbutanoate | p<0.05 | p<0.05 | p<0.05 | p<0.05 | NS | p<0.05 |
| 7 | 3-Methylbutyl acetate | p<0.05 | p<0.05 | p<0.05 | p<0.05 | NS | p<0.05 |
| 8 | Ethyl hexanoate | p<0.05 | p<0.05 | p<0.05 | p<0.05 | NS | p<0.05 |
| 9 | Hexyl acetate | p<0.05 | p<0.05 | NS | p<0.05 | NS | p<0.05 |
| 10 | Ethyl heptanoate | p<0.05 | p<0.05 | NS | NS | NS | p<0.05 |
| 11 | Diethyl succinate | NS | NS | p<0.05 | NS | NS | NS |
|  | **Alcohols** |  |  |  |  |  |  |
| 12 | 2-Mythyl-1-propanol | NS | NS | NS | NS | NS | NS |
| 13 | 1-Butanol | NS | NS | NS | NS | NS | NS |
| 14 | 3Methyl-1-butanol | NS | NS | NS | NS | NS | NS |
| 15 | 1-Hexanol | NS | NS | NS | NS | NS | NS |
| 16 | 3-Ethyl-2-pentanol | p<0.05 | p<0.05 | NS | p<0.05 | p<0.05 | p<0.05 |
| 17 | 2,3-Butanediol | NS | NS | NS | NS | NS | NS |
|  | **Acids** |  |  |  |  |  |  |
| 18 | Acetic acid | p<0.05 | NS | p<0.05 | p<0.05 | p<0.05 | NS |
| 19 | Pentanoic acid | NS | NS | NS | NS | NS | NS |
| 20 | Octanoic acid | p<0.05 | NS | NS | p<0.05 | NS | NS |
|  | **Benzenoid** |  |  |  |  |  |  |
| 21 | 1,3-Ditertiarybutylbenzene | p<0.05 | p<0.05 | NS | p<0.05 | NS | p<0.05 |
|  | **Terpenes** |  |  |  |  |  |  |
| 22 | (-)-Clovene | p<0.05 | p<0.05 | p<0.05 | p<0.05 | p<0.05 | p<0.05 |
| * | **Amylase activity** | NS | NS | NS | NS | NS | NS |

**NS**: Not significant

**Supplementary Table 2** The concentration and odour activity values (OAV) of aroma compounds detected by headspace solid-phase microextraction gas chromatography mass spectrometry (HS-SPME-GC-MS) from the mixture of six types of pooled saliva samples and wine, quantified using external standards.

|  | **Aroma Compound** | **Descriptor** | **Phase A** | | | | **Phase B** | | | | **Phase C** | | | |
| --- | --- | --- | --- | --- | --- | --- | --- | --- | --- | --- | --- | --- | --- | --- |
|  |  |  | **Australian** | | **Chinese** | | **Australian** | | **Chinese** | | **Australian** | | **Chinese** | |
|  |  |  | **Concentration**  **(μg/L)** | **OAV** | **Concentration**  **(μg/L)** | **OAV** | **Concentration**  **(μg/L)** | **OAV** | **Concentration**  **(μg/L)** | **OAV** | **Concentration**  **(μg/L)** | **OAV** | **Concentration**  **(μg/L)** | **OAV** |
|  | **Esters** |  |  |  |  |  |  |  |  |  |  |  |  |  |
| 1 | Ethyl butanoate | Fruity, berry | 452.92 ± 393.49d | 22.65 | 1105.82 ± 28.95b | 55.291 | 1386.07 ± 46.61a | 69.30 | 1370.71 ± 158.4a | 68.54 | 853.36 ± 110.35cd | 42.67 | 1096.17 ± 39.58bc | 54.81 |
| 2 | Ethyl 2-methylbutanoate | Fruity | 157.94 ± 24.71c | 157.94 | 298.98 ± 8.86b | 298.98 | 382.92 ± 16.15a | 382.92 | 381.67 ± 40.58a | 381.67 | 214.21 ± 32.28c | 214.21 | 285.84 ± 8.02b | 285.84 |
| 3 | Ethyl 3-methylbutanoate | Fruity, apple | 209.43 ± 40.39d | 69.81 | 460.45 ± 23.06b | 153.48 | 591.06 ± 11.52a | 197.02 | 578.87 ± 47.14a | 192.96 | 311.36 ± 56.87c | 103.79 | 434.32 ± 21.19b | 144.77 |
| 4 | 3-Methylbutyl acetate | Sweet, fruity, banana | 992.18 ± 146.44d | 4.05 | 1874.03 ± 14.78bc | 7.65 | 2231.6 ± 34.48ab | 9.11 | 2304.3 ± 235.01a | 9.41 | 1352.45 ± 177.72d | 5.52 | 1844.09 ± 97.65c | 7.53 |
| 5 | Methyl hexanoate | Fresh (Pome) | n.d. | / | 21.42 ± 0.75bc | 0.31 | 22.34 ± 0.48ab | 0.32 | 22.96 ± 0.5a | 0.33 | n.d. | / | 20.75 ± 0.33c | 0.30 |
| 6 | Ethyl hexanoate | Apple peel, fruit (Pome) | 756.91 ± 176.14d | 151.38 | 1641.97 ± 41.7bc | 328.39 | 1968.2 ± 52.45ab | 393.64 | 2036.41 ± 198.26a | 407.28 | 1077.92 ± 199.34d | 215.58 | 1526.93 ± 63.65c | 305.39 |
| 7 | Hexyl acetate | Fruity, green | 7.36 ± 4.54c | 0.01 | 12.94 ± 2.05abc | 0.02 | 16.66 ± 0.58a | 0.03 | 17.36 ± 0.82a | 0.03 | 9.44 ± 1.48bc | 0.02 | 13.53 ± 1.23ab | 0.02 |
| 8 | Ethyl heptanoate | Fruity, grape | n.d. | / | 7.88 ± 6.85a | 0.04 | 4.14 ± 7.17a | 0.02 | 12.65 ± 1.11a | 0.06 | n.d. | / | n.d. | 0.00 |
| 9 | Diethyl succinate | Light fruity | 22894.51 ± 1132.07a | 3.82 | 18779.25 ± 1786.05b | 3.13 | 20565.02 ± 1254.85ab | 3.43 | 19256.02 ± 563.85ab | 3.21 | 21276.91 ± 2612.55ab | 3.55 | 22422.04 ± 550.97ab | 3.74 |
|  | **Alcohols** |  |  |  |  |  |  |  |  |  |  |  |  |  |
| 10 | 1-Butanol | Medicinal, alcohol | 2383.76 ± 76.11a | 0.02 | 1493.73 ± 1296.46a | 0.01 | 1717.84 ± 1491.84a | 0.01 | 2488.03 ± 225.85a | 0.02 | 2372.1 ± 67.67a | 0.02 | 2425.55 ± 156.81a | 0.02 |
| 11 | 3-Methyl-1-butanol | Whisky, malt | 505701.39 ± 23544.16a | 16.85 | 452175.44 ± 12399.41a | 15.07 | 510709.08 ± 14899.19a | 17.02 | 481266.72 ± 48552.44a | 16.04 | 486403.25 ± 34110.22a | 16.21 | 505451.47 ± 15100.44a | 16.85 |
| 12 | 1-Hexanol | Green, grass | 4150.9 ± 202.55a | 3.77 | 3969.3 ± 166.38a | 3.61 | 4506.15 ± 172.3a | 4.10 | 4313.7 ± 404.95a | 3.92 | 4094.88 ± 256.39a | 3.72 | 4356.31 ± 171.02a | 3.96 |
|  | **Acids** |  |  |  |  |  |  |  |  |  |  |  |  |  |
| 13 | Pentanoic acid | Rancid, cheesy, sweet | 1449.17 ± 9.18a | 3.73 | 1311.92 ± 75.31a | 3.37 | 1410.46 ± 69.55a | 3.63 | 1411.61 ± 43.91a | 3.63 | 1371.68 ± 56.05a | 3.53 | 1412.83 ± 138.12a | 3.63 |
| 14 | Octanoic acid | Rancid, cheese, fatty acid | 710.55 ± 14.46a | 1.42 | 745.47 ± 37.98a | 1.49 | 777.86 ± 11.7a | 1.56 | 791.59 ± 42.61a | 1.58 | 723.21 ± 26.96a | 1.45 | 770.69 ± 55.53a | 1.54 |
|  | **Benzenoid** |  |  |  |  |  |  |  |  |  |  |  |  |  |
| 15 | Styrene | Plastic | n.d. | / | 5.37 ± 0.11b | - | 6.3 ± 0.2a | - | 6.46 ± 0.38a | - | n.d. | / | 3.33 ± 2.88b | - |
|  | **Terpenes** |  |  |  |  |  |  |  |  |  |  |  |  |  |
| 16 | p-&m-Cymene | Harsh chemical, woody, terpenic | n.d. | / | 0.23 ± 0.4b | - | 1.04 ± 0.08ab | - | 1.26 ± 0.15a | - | n.d. | / | n.d. | / |

*Superscripts letters for the same compound indicate significant differences (*p* < 0.05; Tukey test) on the aroma release from wine after spiking with different saliva samples.

*Odour activity values were calculated by dividing the concentrations by odour threshold ((μg/L) obtained from Guth (1997), Culleré, Escudero, Cacho, and Ferreira (2004), Escudero et al. (2004), Li, Tao, Wang, and Zhang (2008) and Xiao, Zhou, Xiao, and Niu (2017).

*Descriptors are based on the record in the good scents company information system ([http://www.thegoodscentscompany.com](http://www.thegoodscentscompany.com/))

**Supplementary Figure 1 Thirteen species identified as differential among three phases by ANCOM-BC. A)** Point plots with lines showing the log-transformed relative abundance of differential species in three phases. The green lines indicate the species with the lowest relative abundance in Phase A, and the pink lines indicate the species with the lowest relative abundance in Phase B; **B)** Heatmap showing the log-transformed mean relative abundance of differential species.

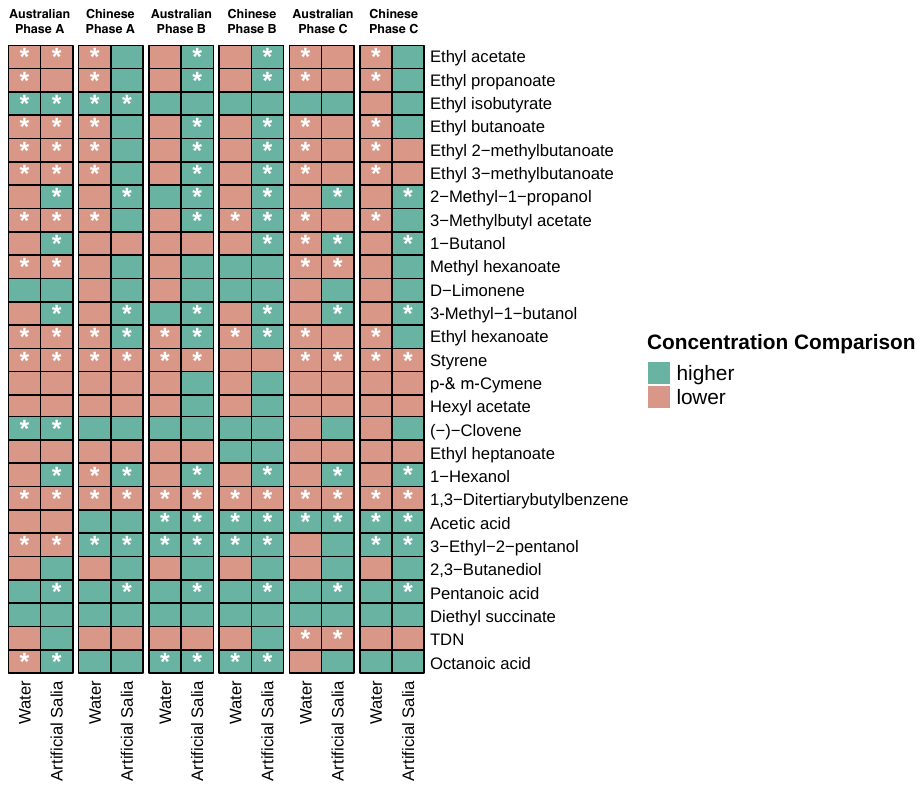
A	B

| **Water** | **Artificial Saliva** |
| --- | --- |
| 9144.88 ± 456.15^a^ | 6490.88 ± 861.03^b^ |
| 92.08 ± 3.87^a^ | 65.59 ± 7.92^b^ |
| 119.61 ± 22.77^a^ | 64.91 ± 2.59^a^ |
| 156.26 ± 6.86^a^ | 114.46 ± 12.35^b^ |
| 105.77 ± 6.92^a^ | 77.14 ± 9.16^b^ |
| 138.57 ± 7.6^a^ | 101.76 ± 15.62^b^ |
| 519 ± 27.88^a^ | 365.14 ± 38.23^b^ |
| 1426.31 ± 83.91^a^ | 1024.6 ± 153.2^b^ |
| 13.8 ± 0.92^a^ | 9.94 ± 1.3^a^ |
| 3.98 ± 0.61^a^ | 1.93 ± 1.77^a^ |
| 0.19 ± 0.33^a^ | n.a. |
| 10571.15 ± 524.32^a^ | 7574.52 ± 941.9^b^ |
| 2529.19 ± 241.38^a^ | 1768.52 ± 359.37^b^ |
| 7.09 ± 1.49^a^ | 5.68 ± 0.83^a^ |
| 6.98 ± 7.97^a^ | 4.91± 2.29^a^ |
| 22.18 ± 7.09^a^ | 16.93 ± 4.32^a^ |
| 28.74 ± 47.57^a^ | 10.32 ± 4.65^a^ |
| 5.08 ± 7.51^a^ | 4.44 ± 3.42^a^ |
| 351.24 ± 19.14^a^ | 248.03 ± 35.02^b^ |
| 185.33 ± 48.61^a^ | 136.93 ± 31.7^a^ |
| 147.75 ± 38.4^a^ | 108.52 ± 7.83^a^ |
| 30.37 ± 2.66^a^ | 24.31 ± 2.94^a^ |
| 52.06 ± 19.73^a^ | 30.12 ± 5.52^a^ |
| 7.26 ± 1.18^a^ | 3.38 ± 1.03^b^ |
| 1178.12 ± 1020.34^a^ | 1857.49 ± 466.16^a^ |
| 29.34 ± 15.96^a^ | 15.97 ± 3.68^a^ |
| 5.24 ± 0.38^a^ | 3.3 ± 0.8^b^ |

**Supplementary Figure 2 The aroma released from wine after mixing with water, artificial saliva, and pooled saliva samples. A)** The table showing the concentration of volatiles released from wine-water and wine-artificial saliva mixture. Superscripts indicate significant differences (p < 0.05; Student’s t test) between the two groups. **B)** The coloured squares indicate the comparison among wine-water, wine-artificial saliva, and wine-saliva mixtures. The green squares represent the higher concentration of released aroma after mixing with the pooled saliva than mixing with water or artificial saliva. The pink squares represent the lower concentration of released aroma after mixing with the pooled saliva than mixing with water or artificial saliva. The grey squares represent the volatiles that have not been detected. The significant differences (p < 0.05; one-way ANOVA) were indicated by “*”.

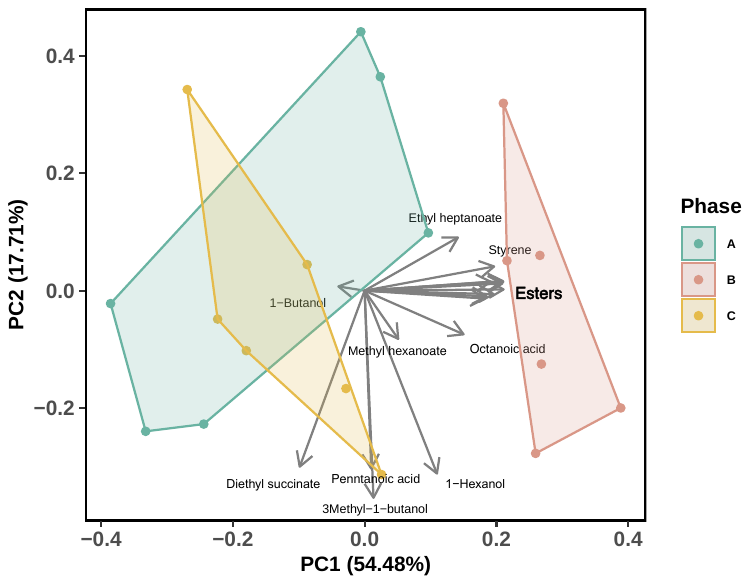

A	B

**Supplementary Figure 3 Principal component analysis (PCA) plots of A) 27 semi-quantified compounds and B) 17 compounds quantified using external standards showing the similar results with the PCA plots of 21 semi-quantified compounds detected constantly in all subgroups.**
